# *Tcf4* encodescortical differentiation during development

**DOI:** 10.1101/470385

**Authors:** Simone Mesman, Reinier Bakker, Marten P. Smidt

**Affiliations:** Swammerdam Institute for Life Sciences FNWI University of Amsterdam, Science Park 904 1098XH Amsterdam, The Netherlands

**Keywords:** Pitt-Hopkins Syndrome, *Tcf4*, cortical development, layering deficit, human brain defects, E-box factors

## Abstract

*Tcf4* has been linked to autism, schizophrenia, and Pitt-Hopkins Syndrome (PTHS) in humans, however, the mechanisms behind its role in disease development is still elusive. In the present study, we provide evidence that *Tcf4* has a critical function in the differentiation of cortical regions during development.

We show that *Tcf4* is present throughout the developing brain at the peak of neurogenesis. Deletion of *Tcf4* results in mis-specification of the cortical layers, malformation of the corpus callosum and hypoplasia of the hippocampus. RNA-sequencing on E14.5 cortex material shows that *Tcf4* functions as a transcriptional activator and loss of *Tcf4* results in downregulation of genes linked to the emergence of other neurodevelopmental disorders. Taken together, we show that neurogenesis and differentiation are severely affected in *Tcf4* mutants, phenocopying morphological brain defects detected in PTHS patients. The presented data identifies new leads to understand the mechanism of human brain defects and will assist in genetic counseling programs.

## 1. Introduction

Correct cortical neurogenesis depends on a complex genetic program executed through correct spatio-temporal expression of transcription factors. A specific group of transcription factors present in the developing cortex, is the basic helix-loop-helix (bHLH) protein family (Powell and Jarman, 2008). The E-box protein sub-family of bHLH proteins consists out of three members; *Tcf3* (*E2A*), *Tcf12* (*HEB*), and *Tcf4* (*E2-2*), which functions are dependent on specific bHLH binding partners through homo- or heterodimerization (Massari and Murre, 2000; Murre et al., 1989; Powell and Jarman, 2008). Mutations in the bHLH-domain, comprising the DNA-binding interface, inhibits such dimerization processes and proper DNA-binding, which directly interferes with their function in gene regulatory events (Sweatt, 2013). E-box proteins have mainly been studied in relation to their role in immune system development. They are found to be critical for the transition from CD4^+^CD8^+^ double-positive T-cells to CD4^+^ or CD8^+^ single-positive T-cells (Wojciechowski et al., 2007) and loss of either *Tcf12* or *Tcf3* leads to an early depletion of T-cell progenitors and a decrease in mature T-cells (Wojciechowski et al., 2007). *Tcf4* on the other hand, has only a minor role during early thymocyte development and it has been suggested that compensatory mechanism through *Tcf3* and/or *Tcf12* exists (Wikström et al., 2008).

In the brain, E-box proteins are thought to regulate neurogenesis and neuronal differentiation based on their spatio-temporal presence and binding partners (Powell and Jarman, 2008). In humans, haplo-insufficiency of *Tcf4* has been found to be determinative for Pitt-Hopkins Syndrome (PTHS) (Brockschmidt et al., 2007; Marangi et al., 2011; Marangi et al., 2012; Sweatt, 2013), a rare mental disorder, hallmarked by severe intellectual disability, typical facial gestalt, and additional features like breathing problems (Blake et al., 2010; Hasi et al., 2011; Peippo and Ignatius, 2012). Imaging of PTHS patients brains shows major neurodevelopmental defects, ranging from a smaller corpus callosum, underdeveloped hippocampi, and defective cortical development, to microcephaly, enlarged ventricles, and bulging caudate nuclei (Blake et al., 2010; Ghosh et al., 2012; Hasi et al., 2011; Peippo and Ignatius, 2012). Finally, *Tcf4* has been implicated in other developmental brain disorders, like schizophrenia and autism (Brzózka and Rossner, 2013; Cousijn et al., 2014; Wirgenes et al., 2012).

Until now the mechanism behind the observed neurodevelopmental defects as observed in humans is not known. Previous studies on a *Tcf4* mouse mutant describe a loss of the pontine nucleus (Flora et al., 2007). Furthermore, Fischer et al., 2014 reported that *Tcf4* promotes differentiation of neural stem cells to neurons in the adult forebrain and may thus be involved in postnatal neurogenesis by orchestrating neural stem cell lineage progression. In mammalian cell-lines *Tcf4* has been found to interact with *Mash1* and *NeuroD2*, important regulators of neuronal differentiation (Persson et al., 2000). Studies by Jung et al. (2018) and Grubišić et al. (2015) showed that the phenotype detected in the brain and in the gut of patients with PTHS could also (partially) be detected in heterozygous mutants of TCF4. Taken together, the available data point to a possible function of *Tcf4* in neuronal development and links the developmental defects detected in heterozygous mutants to the phenotype in PTHS patients. In this study we aimed to gain more insight in and further specify the function of *Tcf4* in the developing cortex. We demonstrate the specific presence of *Tcf4* in specific cortical layers during development. Ablation of *Tcf4* (Zhuang et al., 1996) induces major defects in cortical development, as cortical layering and specification of layer-specific neurons is severely affected. Furthermore, CUX1 positive neurons, a signature for upper layer neurons, are absent in *Tcf4* mutants, whereas CTIP2 positive neurons (lower layer neurons) are decreased and show aberrant co-localization with SATB2. Besides these differentiation defects, the corpus callosum and both hippocampi are severely underdeveloped in these mutants. Transcriptome analyzes through RNA-sequencing on E14.5 cortical material indicate that *Tcf4* is an activator of gene expression in the developing cortex and is activates genes that are linked to neurogenesis and neuronal maturation. Importantly, *Tcf4* regulates genes that are known for their involvement in clinical syndromes related to PTHS and ID. Finally, the observed brain malformations phenocopy clinical features observed in PTHS patients, further establishing this mouse mutant as an excellent model for this human developmental disorder, providing mechanistic insight to the syndrome and assist in the identification of novel factors for genetic screening and genetic counseling for human patients with neurodevelopmental disorders.

## 2. Materials and Methods

### 2.1 Ethics Statement

All animal studies are performed in accordance with local animal welfare regulations, as this project has been approved by the animal experimental committee (Dier ethische commissie, Universiteit van Amsterdam; DEC-UvA), and international guidelines

### 2.2 Animals

The transgenic mouse line B6;129-Tcf4^tm1Zhu^/J originates from the Jackson Laboratory. B6;129-Tcf4^tm1Zhu^/J mouse line was back-crossed with the C57BL/6 line and knock-out embryos were generated by crossing heterozygous *Tcf4* mice. WT embryos, to examine WT expression patterns of *Tcf4*, were generated by crossing C57BL/6 mice. Pregnant mice [embryonic day 0.5 (E0.5) is defined as the morning of plug formation] were sacrificed by cervical dislocation. Embryos were collected in 1x PBS and immediately frozen on dry-ice, or fixed by immersion of 3-12 h in 4% paraformaldehyde (PFA) at 4°C. After PFA incubation samples were washed in 1x PBS and cryoprotected O/N at 4°C in 30% sucrose. Embryos were frozen on dry-ice and stored at −80°C. Cryosections were cut at 16 μm, mounted on Superfrost Plus slides (Thermo Fisher Scientific), air-dried, and stored at −80°C until further use.

### 2.3 Genotyping

Genotyping of B6;129-Tcf4^tm1Zhu^/J transgenic embryos and mice was performed according to the protocol of the Jackson Laboratory. Briefly, 100 ng of genomic DNA was used together with primer pair: FP 5’-AGCGCGAGAAAGGAACGGAGGA-3’, RP1 5’-GGCAATTCTCGGGAGGGTGCTT-3’, and RP2 5’-CCAGAAAGCGAAGGAGCA-3’ resulting in a product at 229 bp for the WT allele and a product at 400 bp for the KO allele.

### 2.4 *In situ* hybridization

*In situ* hybridization with digoxigen (DIG)-labeled probes was performed as described previously (Smidt et al., 2004). Fresh frozen sections were fixed in 4% PFA for 30 min and acetylated with 0.25% acetic anhydride in 0.1 M triethanolamine for 10 min. Probe hybridization was carried out at 68°C O/N with a probe concentration of 0.4 ng/μl in a hybridization solution containing 50% deionized formamide, 5x SSC, 5x Denhardt’s solution, 250 μg/mL tRNA Baker’s yeast, and 500 μg/mL sonificated salmon sperm DNA. The following day slides were washed in 0.2x SSC for 2 h at 68°C followed by blocking with 10% heat-inactivated fetal calf serum (HIFCS) in buffer 1 (100 mM Tris Hcl, pH 7.4 and 150 mM NaCl) for 1 h at RT. DIG-labeled probes were detected by incubating with an alkaline-phosphatase-labeled anti-DIG antibody (Roche, Mannheim, 1:5000), using NBT-BCIP as a substrate. Slides were washed 2×5 min in T_10_E_5_, dehydrated with ethanol and embedded in Entellan.

DIG *in situ* hybridization was performed with a 918 bp *Tcf4* fragment bp 1101-2018 of mouse cDNA.

### 2.5 Immunohistochemistry

Fluorescence immunohistochemistry (IHC) was carried out as described previously (Fenstermaker et al., 2010; Kolk et al., 2009). Cryosections were blocked with 4% HIFCS in 1x THZT and incubated with a primary antibody [Rb-Gap43 (ab16053 Abcam 1:1000), Ms-Tcf4 (H00006925-M03 Abnova 1:500. Antibody specificity has been shown by Forrest et al. (2013) and Tanaka et al. (2010)), Rb-Ctip2 (ab28488 Abcam, 1:1000), Ms-Satb2 (ab51502 Abcam, 1:1000), Rb-Tbr1 (ab31940 Abcam, 1:1000), Rb-Tbr2 (ab23345 Abcam, 1:1000), Rb-Ki67 (ab15580 Abcam 1:500), Rb-Cux1 (sc-13024 Santa Cruz, 1:100)] diluted in 1x THZT O/N at RT. The next day sections were incubated with a secondary Alexafluor antibody (anti-rabbit, anti-mouse) diluted 1:1000 in 1x TBS for 2 h at RT. This procedure was repeated for double-labeling with a different primary antibody. After immunostaining nuclei were stained with DAPI (1:3000) and washed extensively in 1x PBS. Slides were embedded in Fluorsave (Calbiogen) and analyzed with the use of a fluorescent microscope (Leica). Antibodies against TCF4, CTIP2, SATB2, TBR1, and TBR2 required antigen retrieval as follows. Slides were incubated with 0.1 M citrate buffer pH6 for 3 min at 800 W and 9 min a 400 W, cooled down to RT in a water bath, after which the protocol was followed as usual.

Quantification of Ki67-, CTIP2-, and SATB2-expressing cells was performed in 4-7 (matching) coronal sections (WT n=3-4;KO n=3-4). To determine the cellular position of CTIP2-expressing cells, a grid containing 10 bins was placed over coronal sections before quantification. Cells were counted as CTIP2^+^ or Ki67^+^ when staining co-localized with the nuclear DAPI staining, and as double positive when staining of CTIP2 and SATB2 co-localized. Statistical analysis was performed via a one- or two-tailed student’s t-test.

### 2.6 RNA-sequencing

RNA was isolated from dissected E14.5 cortices of *Tcf4* WT and *Tcf4* mutant embryos. RNA was isolated with Trizol (ThermoFisher) according to manufacturers protocol. After isolation RNA clean-up was performed with an RNA mini kit from Qiagen according to manufacturers protocol. Isolated RNA of dissected cortices of 2 WT or mutant embryos were pooled to eventually form an n=3. Pair-end RNA-sequencing (minimal 2* 10^6 reads per sample), mapping on the mouse genome and statistical analysis on read counts was performed commercially (Service XS, Leiden, The Netherlands).

GO-term analysis was performed via the PANTHER over-representation test (release 20150430) with Bonferroni correction for multiple comparisons.

## 3. Results

### 3.1 *Tcf4* is present in cortical neurons at different stages of development

In order to determine the role of *Tcf4* in cortical development we first assessed its spatio-temporal expression pattern (**Figure 1A**). At E14.5 *Tcf4* transcript is detected throughout the developing brain, although it is most abundantly expressed in the cortex. At E16.5 the expression of *Tcf4* is still present throughout the cortex, however in the ventricular - and subventricular zone and outer parts of the cortical plate transcript levels appear to be higher, suggesting a layer specific intensity of the *Tcf4* transcript.

**Figure 1:**
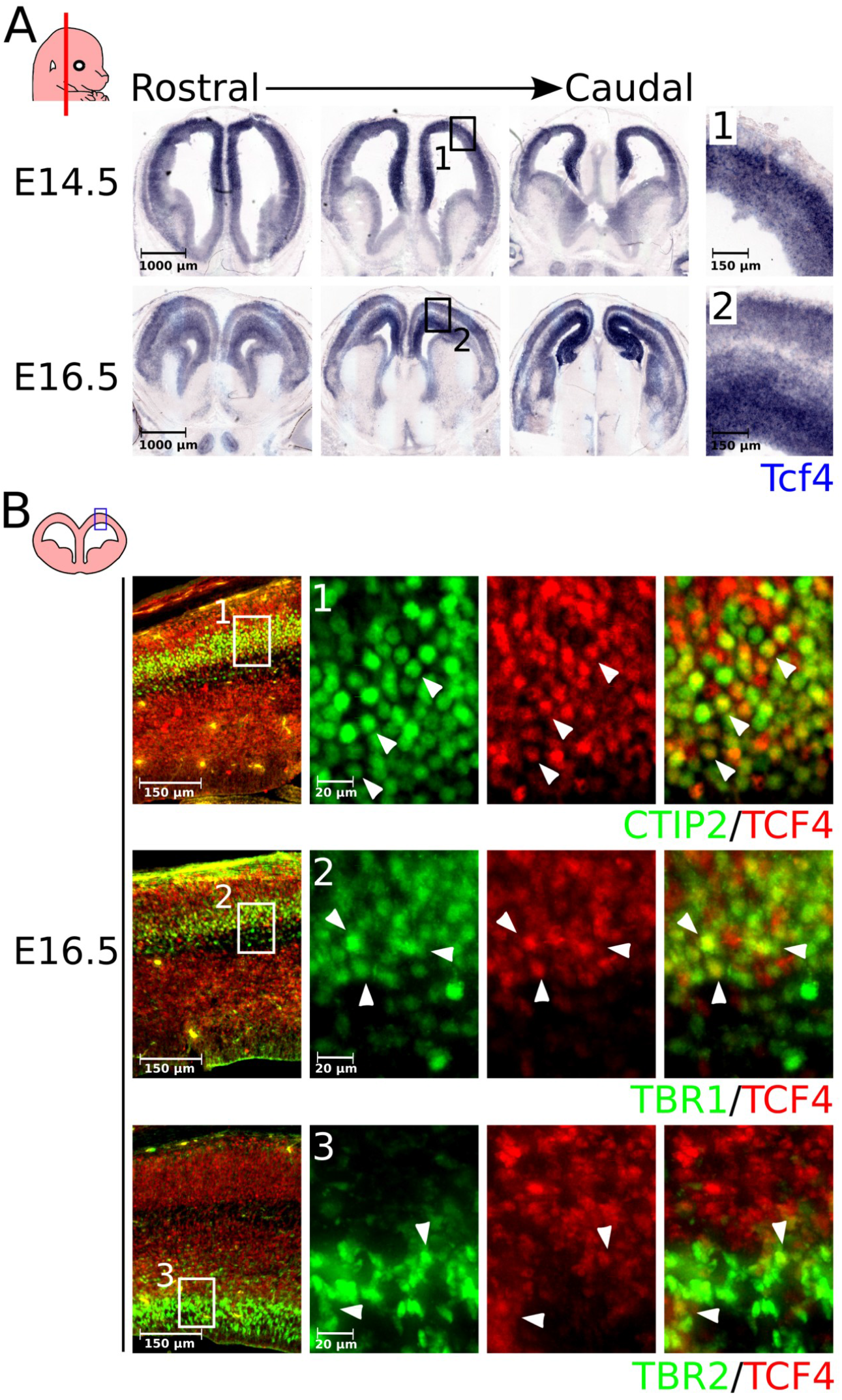
*Tcf4* is expressed throughout the mouse brain during development and has a layer-specific expression in the cortex. (**A**) *Tcf4* (blue) is expressed throughout the mouse brain at E14.5 and E16.5. At El4.5 *Tcf4* expression is most prominent in the cortex, where it is expressed throughout the structure (**1**). At El6.5 expression of *Tcf4* is still present in the cortex and becomes even further specified in the different cortical layers (**2**). (**B**) TCF4 protein is expressed in a layer-specific fashion at El6.5. TCF4 (red) protein expression shows strong co-localization with the layer specific markers CTIP2 (green) (**1**, white arrowheads) and TBR1 (green) (**2**, white arrowheads). However, TCF4 only slightly overlaps with the TBR2^+^ (green) neural progenitor population of the VZ and SVZ (**3**, white arrowheads).

To confirm these data and to determine in which cortical layers TCF4 protein is present, we examined the cortex during development. As TCF4 protein was not detected in the cortex until E14.5 and becomes more apparent at later stages (**Supplemental Figure 1**), we focused on the expression pattern of TCF4 and layer-specific marks at E16.5 when most neurons of the developing cortex are specified (Molyneaux et al., 2007). At E16.5, similar to *Tcf4* transcript, TCF4 is found in specific layers of the developing cortex. It co-localizes with CTIP2 (**Figure 1B-1** white arrowheads), a marker for layer V and VI, and TBR1 (**Figure 1B-2** white arrowheads), a marker for layer VI. Although *Tcf4* transcript was detected in the ventricular- and subventricular zone at this stage, TCF4 protein is only marginally present in the VZ and SVZ of the cortex, identified by TBR2. Finally, some TBR2-positive cells, bordering the intermediate zone of the cortex, co-localize with TCF4 (**Figure 1B-3** white arrowheads).

### 3.2 Loss of *Tcf4* results in defective cortical layering

As shown above, *Tcf4* is present in the murine developing cortex at the peek of neurogenesis. In order to determine the consequence of heterozygous or full deletion of *Tcf4* on development of the cortex, we first examined the general cortical structure by means of DAPI-staining. At E17.5 the WT cortex shows a clear distinction in different layers. Based on neuronal density we were able to divide the cortex into a cortical plate (CP), Layer VI, intermediate zone (IZ), and the ventricular zone/subventricular zone ((S)VZ). Interestingly, the clear distinction between the IZ and layer VI was not present in *Tcf4* heterozygous and full mutant embryos (**Figure 2A** white asterisk). Furthermore, the overall cortical thickness (CT) is reduced with ~14% in full mutants compared to WT (n=4; p<0.05, one-tailed), but is not significantly different between heterozygous and mutant or WT embryos (**Figure 2B**). Besides the decrease in CT, a decrease in cortical plate thickness (CP) was detected in the full mutant of ~29% compared to WT (n=4; p<0.001, two-tailed) and ~20% compared to heterozygous embryos (n=4; p<0.01, two-tailed) (**Figure 2C**). This change is reflected in the ratio of the CP compared to the CT, which is ~6% smaller in full mutants compared to WT (n=4; p<0.01, two-tailed) and ~4% smaller compared to heterozygous animals (n=4; p<0.05, two-tailed) (**Figure 2D**).

**Figure 2:**
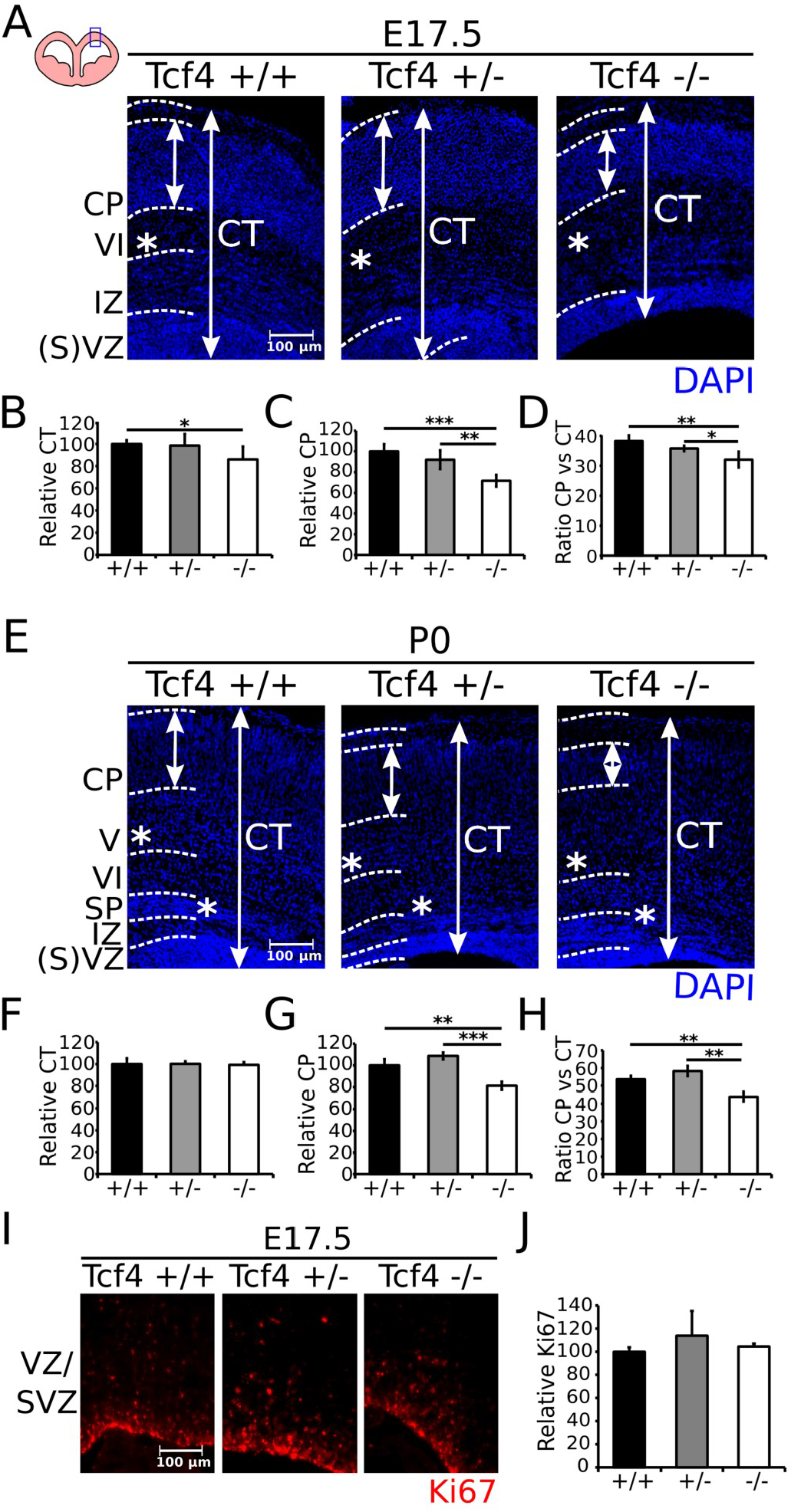
*Tcf4* deletion leads to a general effect on the cortical anatomy at E17.5 and P0. (**A**) Compartmentalization of the cortex at El7.5 is affected in the *Tcf4* heterozygous and mutant embryos at El7.5. DAPI staining (blue) shows a less prominent distinction of the cortical layers in the heterozygous and mutant cortex (white asterisk). (**B**) Cortical thickness (CT) is decreased with −14% in the mutant (white bars) compared to the WT (black bars) (n=4; * p<0.05, one-tailed), but is not altered in the heterozygous embryos (gray bars). WT was set at 100%. (**C**) The cortical plate (CP) is −29% smaller in the mutant compared to the WT (n=4; *** p<0.001, two-tailed) and −21% smaller compared to heterozygous embryos (n=4; ** p<0.01, two-tailed). WT was set at 100%. (**D**) A similar shift was seen in the ratio of CP and CT, which is −6% smaller in the mutant compared to the WT (n=4; ** p<0.01, two-tailed) and −4% smaller compared to heterozygous embryos (n=4; * p<0.05, two-tailed). WT was set at 100%. (**E**) Compartmentalization of the cortex remains affected at P0 in the heterozygous and mutant animals. DAPI staining (blue) shows a less prominent distinction of the cortical layers in the heterozygous and mutant cortex (white asterisks). (**F**) Cortical thickness (CT) is no longer affected at P0 between the WT (black bars), heterozygous (gray bars), and mutant (white bars) animals. WT was set at 100%. (**G**) The thickness of the cortical plate (CP) is decreased with ~19% in the mutant compared to the WT (n=3; ** p<0.01, two-tailed) and with ~28% compared to heterozygous animals (n=3; *** p<0.001, two-tailed). WT was set at 100%. (**H**) A similar shift was seen in the ratio between the CP and the entire cortex, which is ~10% smaller in the mutant compared to the WT (n=3; ** p<0.01, two-tailed) and ~15% smaller compared to heterozygous animals (n-3; ** p<0.01, two-tailed). WT was set at 100%. (**I**) Loss of *Tcf4* does not affect the proliferation of stem cells in the VZ and SVZ of the developing cortex at E17.5. The proliferation mark Ki67 (red) was normally present in the WT, heterozygous, and mutant cortex. (**J**) Quantification shows no difference in the amount of Ki67^+^ cells at E17.5 between WT (black bar), heterozygous (gray bar), and mutant (white bar) animals (n=3). WT was set at 100%

At P0 we were able to distinguish several cortical layers based on cell-density; CP, Layer V-VI, sub-plate (SP), IZ, and VZ and SVZ. Similar as observed at E17.5, at P0 the layers of the cortex are less distinctive in both the heterozygous and mutant animals (**Figure 2E** white asterisks), although the CT area has been recovered (**Figure 2F**). The thickness of the CP is ~19% smaller in mutants compared to WT (n=3; p<0.01, two-tailed), and ~27% smaller compared to heterozygous animals (n=3; p<0.001, two-tailed) (**Figure 2G**). Notably, the CP of the heterozygous animals shows an upward trend in thickness, as this is ~9% increased compared to WT animals (n=3; p=0.08, two-tailed) at P0. The ratio between the CP and the CT is more severely affected at P0 (**Figure 2H**). The ratio of the CP to the CT is ~54% in WT and ~58% in the heterozygous animals, which is decreased to ~44% in mutant animals compared to WT (n=3; p=<0.01, two-tailed) and heterozygous (n=3; p<0.01, two-tailed) animals.

Although TCF4 is not clearly present in the VZ and SVZ of the developing cortex, the increase in CT of the mutant cortex between E17.5 and P0 could be due to an elevated proliferation at E17.5 (**Figure 2I**). Ki67 labeling, marking proliferating cells, showed that there is no significant difference in proliferation in the cortex of mutant or heterozygous embryos at E17.5 compared to WT (n=3; two-tailed) (**Figure 2J**). Together, these data indicate that the correct developmental process of the murine cortex is affected upon loss of one or two alleles of *Tcf4*.

### 3.3 Loss of *Tcf4* results in mis-specification of CTIP2- and SATB2-expressing neurons at E17.5

Above we have shown that loss of one or two alleles of *Tcf4* results in a difference in neuronal distribution throughout the cortex, suggesting a defective cortical layering at E17.5 and P0. To determine whether expression of layer-specific markers in the cortex is affected accordingly, we performed immunohistochemistry for SATB2 and CTIP2 at E17.5 in *Tcf4* mutants (-/- and +/-) compared to WT. These two proteins mark two different neuronal populations, as SATB2^+^ neurons are callosal projection neurons (Alcamo et al., 2008; Leone et al., 2015; Srinivasan et al., 2012; Srivatsa et al., 2014) and CTIP2^+^ neurons project to other cortical areas in the same hemisphere (Leid et al., 2004; Srinivasan et al., 2012; Srivatsa et al., 2014). SATB2 is present throughout the differentiated layers of the cortex and not in the VZ and SVZ in *Tcf4* WT, heterozygous, and mutant cortices at E17.5 (**Figure 3A**). CTIP2, a marker for neurons in layer V and VI of the cortex, is also present in *Tcf4* heterozygous and mutant cortices (**Figure 3B**). However, quantification of CTIP2^+^ neurons at E17.5 shows that in full mutants there is a significant decrease of ~20% and ~17% in the amount of CTIP2^+^ cells compared to WT (n=4; p<0.05, one-tailed) and heterozygous (n=4; p<0.01, one-tailed) animals, respectively (**Figure 3D**).

**Figure 3:**
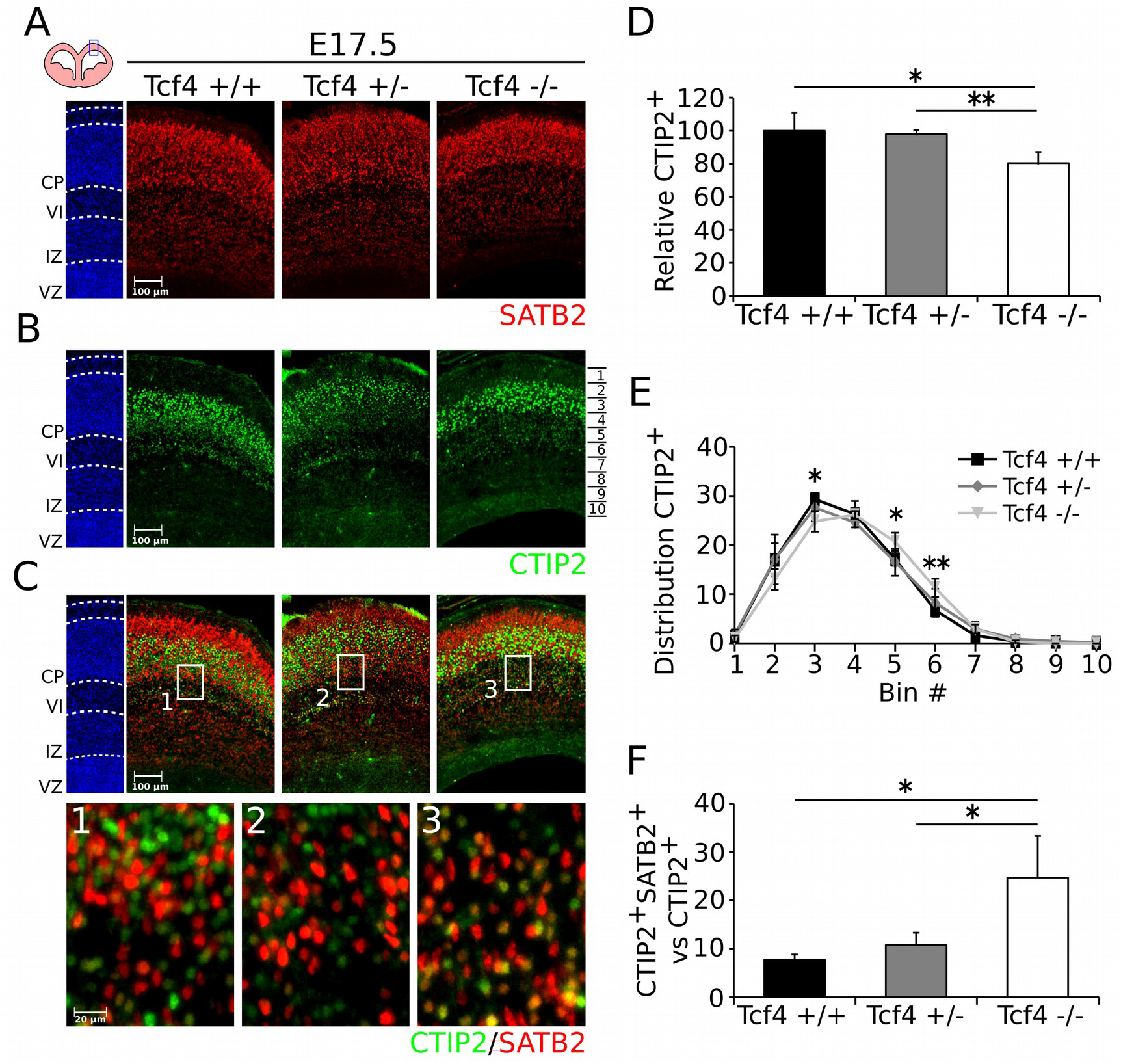
Deletion of *Tcf4* results in an aberrant distribution and mis-specification of CTIP2^+^ cells in the cortex at E17.5. (**A**) SATB2 (red) expression is normally present in the heterozygous and mutant cortex compared to the WT at E17.5. (**B**) Expression of CTIP2 (green) is present in the heterozygous and mutant cortex at E17.5, although the distribution of CTIP2-expressing cells appears to be altered in the mutant compared to the WT cortex. (**C**) Co-localization of SATB2 (red) and CTIP2 (green) is more apparent in the mutant cortex (**3**) compared to the WT (**1**) and heterozygous (**2**) cortex at E17.5. (**D**) Expression of CTIP2 is decreased with ~20% in the mutant compared to the WT (n=3; * p<0.05, one-tailed) and ~17% compared to the heterozygous (n=3; ** p<0.01, one-tailed) embryos. WT was set at 100%. (**E**) Distribution of CTIP2^+^ cells is slightly shifted to lower parts of the cortex in the mutant (light gray line) compared to WT (black line), but not to heterozygous embryos (gray line), bin 3 ~5% decrease (n=4; * p<0.05, two-tailed), bin 5 ~3% increase (n=4; * p<0.05, two-tailed), and bin 6 ~4.5% increase (n=4; ** p<0.01, two-tailed) between mutant and WT embryos. Ratio to total CTIP2^+^ cells. (**F**) The amount of CTIP2^+^/SATB2^+^ neurons in the CTIP2^+^ population shows an increase of ~17% between mutant (white bar) and WT (black bar) embryos (n=3; * p<0.05, one-tailed), and of ~14% between the mutant and heterozygous (gray bar) embryos (n=3; * p<0.05, one-tailed).

To determine the distribution of CTIP2-expressing cells in the cortex, we divided the cortex in 10 bins (bin 1 positioned in the upper part of the cortex to bin 10 at the VZ of the cortex) and quantified the CTIP2-expressing cells per bin (**Fig3B** and **E**). We detected a slight shift of CTIP2^+^ cells to lower bins in the *Tcf4* mutant compared to the WT. In bin 3 less CTIP2^+^ cells were observed (~29% of total in the WT compared to ~24.8% of total in the mutant) (n=4, p<0.05, two-tailed), whereas more CTIP2^+^ cells were detected in bin 5 (WT ~17%, mutant ~20.7%; n=4; p<0.05, two-tailed), and 6 (WT ~6.7%, mutant ~11.3%; n=4; p<0.01, two-tailed). No difference in distribution was detected between WT and heterozygous embryos at E17.5, although an increase in amount of CTIP2^+^ cells was detected between the heterozygous and mutant embryos in bin 5 (heterozygous ~16.5%, mutant ~20.7%; n=4; p<0.05, two-tailed).

Since the distribution of CTIP2^+^ cells is altered in *Tcf4* mutants, we set out to determine whether the identity of these neurons is changed accordingly. In wild type animals ~5% of the total amount of CTIP2^+^ cells in layer VI co-localizes with SATB2 cells (Srinivasan et al., 2012). Since we described a shift of CTIP2 cells towards lower parts, we determined whether the SATB2/CTIP2 co-localizing population may be over-represented in *Tcf4* mutants (**Figure 3C**). In the cortex of WT embryos at E17.5 we found that ~7.3% of the total CTIP2^+^ population co-localizes with SATB2-expressing neurons (**Figure 3C-1** and **F**), similar to what was described earlier (Srinivasan et al., 2012). In heterozygous embryos (**Figure 3C-2**) an upward trend was observed in the amount of double-positive cells towards ~10.8% (n=4; p=0.06, one-tailed). Interestingly, this apparent increase became significantly higher in full *Tcf4* mutants (~24.7%, **Figure 3C-3**) compared to WT (n=4; p<0.05, one-tailed) and heterozygous (n=4; p<0.05, one-tailed) animals (**Figure 3F**).

### 3.4 Deletion of *Tcf4* results in a mis-specification of CTIP2- and SATB2-expressing neurons and loss of the Cux1^+^ neuronal population at P0

Above we have shown that loss of one or two alleles of *Tcf4* results in a defective segregation of CTIP2 and SATB2 neurons in the cerebral cortex. To determine whether this phenotype is persistent after birth, we performed similar experiments on P0 animals. Expression of SATB2 is still relatively normal at P0 in the heterozygous and mutant cortex compared to the WT (**Figure 4A**). The amount of CTIP2-expressing cells on the other hand still displays a significant decrease (**Figure 4B**), although this decrease was lowered to ~12% between the mutant and the WT (n=3; p<0.05, one-tailed) and no significant difference was detected anymore between heterozygous and full mutants (**Figure 4D**). Interestingly, the shift in distribution to lower parts of the cortex was still increased at P0 between WT and mutant animals, and a similar shift was detected for heterozygous compared to WT animals (**Figure 4E**). Bin 4 (WT ~21.6%, heterozygous ~13.9%, mutant ~14.7%; n=3; p<0.001, two-tailed) and 5 (WT ~22%, heterozygous ~18%, mutant ~15.3%; n=3; p<0.001, two-tailed) showed a decrease in CTIP2^+^ neurons for both mutant and heterozygous compared to WT animals, whereas bin 7 (WT ~11%, heterozygous ~18%, mutant ~21.5%; n=3; p<0.001, p<0.01 respectively, two-tailed), bin 8 (WT ~2.6%, heterozygous ~7.8%, mutant ~9.3%; n=3; p<0.05, two-tailed), and bin 9 (WT ~0.7%, heterozygous ~2.9%, mutant ~1.8%; n=3; p<0.05, n.s. respectively, two-tailed) showed an increase in the amount of CTIP2-expressing cells compared to WT animals (**Figure 4E**).

**Figure 4:**
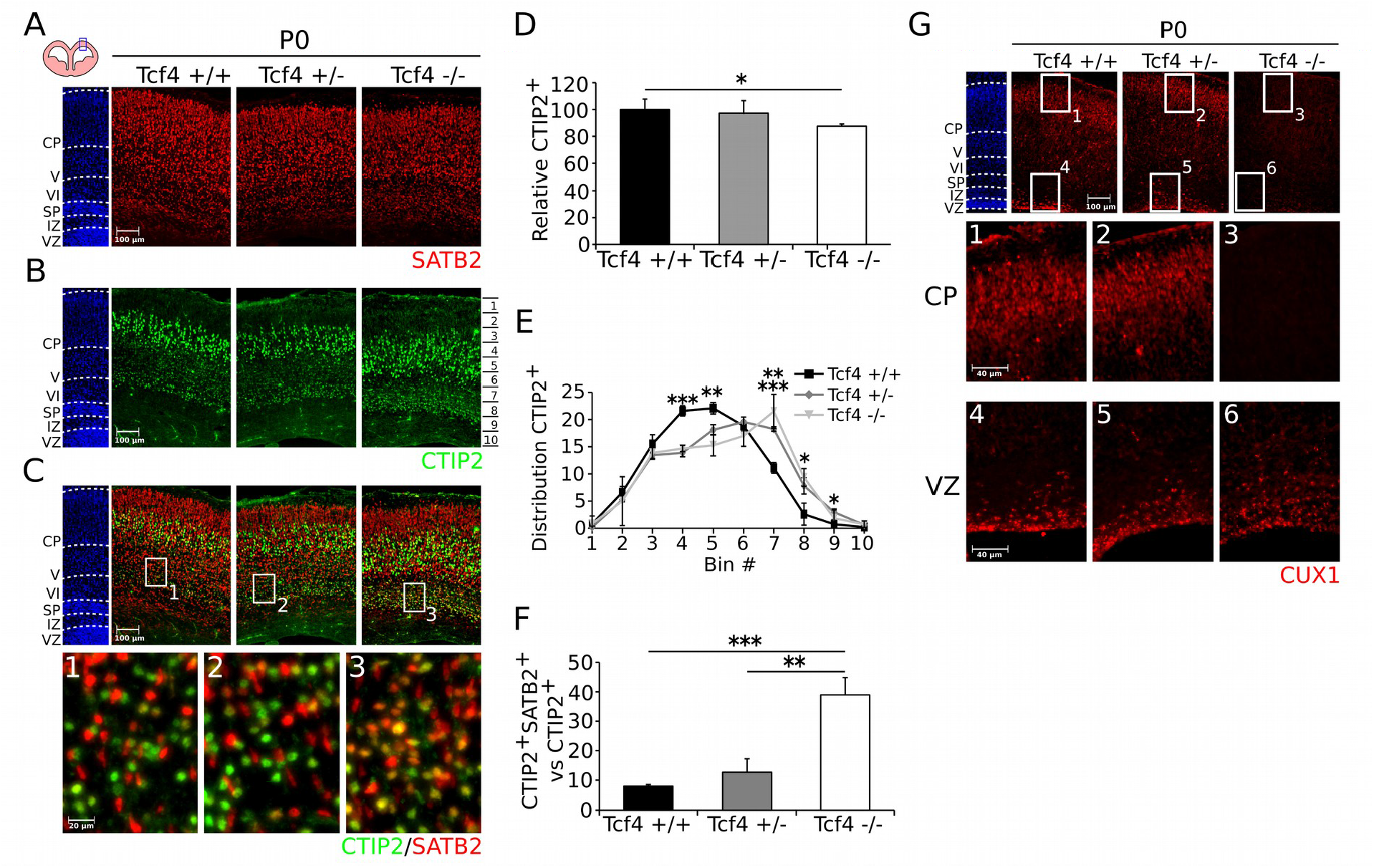
Deletion of *Tcf4* results in an aberrant distribution and mis-specification of CTIP2^+^ cells and a loss of CUX1-expressing cells at P0. (**A**) SATB2-expression (red) shows a normal expression in the heterozygous and mutant cortex at P0. (**B**) CTIP2^+^ cells (green) show a change in distribution in the heterozygous and mutant cortex compared to the WT at P0. (**C**) Co-localization of SATB2 (red) and CTIP2 (green) is more apparent in the mutant cortex (**3**) compared to the WT (**1**) and heterozygous (**2**) cortex at P0. (**D**) The amount of CTIP2^+^ cells is decreased with ~12% in the mutant (white bar) compared to the WT (black bar) (n=3; * p<0.05, one-tailed), but not compared to the heterozygous animals (gray bar). WT was set at 100%. (**E**) The distribution of CTIP2^+^ cells shows a shift to lower parts of the cortex in the mutant (light gray line) compared to the WT (black line), bin 4 ~7% decrease (n=3; *** p<0.001, two-tailed), bin 5 ~7% decrease (n=3; ** p<0.01, two-tailed), bin 7 ~10% increase (n=3; ** p<0.01, two-tailed), bin 8 ~7% increase (n=3; * p<0.05, two-tailed) between mutant and WT animals. Heterozygous animals (gray line) show a similar shift compared to the WT, bin 1 ~0.6% decrease (n=3; * p<0.05, two-tailed), bin 4 ~8% decrease (n=3; *** p<0.001, two-tailed), bin 5 ~4% decrease (n=3; ** p<0.01, two-tailed), bin 7 ~7% increase (n=3; *** p<0.001, two-tailed), bin 8 ~5% increase (n=3; * p<0.05, two-tailed), and bin 9 ~1.2% increase (n=3; * p<0.05, two-tailed) between WT and heterozygous animals. Ratio to total CTIP2^+^ cells. (**F**) The amount of CTIP2^+^/SATB2^+^ neurons in the CTIP2^+^ population shows an increase of ~31% between mutant (white bar) and WT (black bar) embryos (n=3; *** p<0.001, one-tailed), and of ~26% between mutant and heterozygous (gray bar) embryos (n=3; ** p<0.01, one-tailed). (**G**) CUX1 is normally expressed in the WT and heterozygous animals at P0, but completely lost in the CP of the mutant cortex (**1-3**). CUX1 expression in the VZ is still present in WT, heterozygous, and mutant cortex (**4-6**).

Quantification of the population CTIP2^+^/SATB2^+^ neurons showed that, confirming the E17.5 data, the amount of double-positive cells was ~8% of the CTIP2^+^ population in the WT cortex (**Figure 4C-1** and **F**). The amount of CTIP2^+^/SATB2^+^ neurons showed an upward trend to ~12.7% in heterozygous animals (**Figure 4C-2)** compared to WT animals (n=3; p=0.08, one-tailed). Finally, the amount of double-positive neurons in the mutant cortex (**Figure 4C-3**) increased to ~39% of the total CTIP2-expressing population, which was significant compared to both WT (n=3; p<0.001, one-tailed) and heterozygous (n=3; p<0.01, one-tailed) animals (**Figure 4F**).

As the mis-specification of the cortical layers was more severely affected at P0 and the CP displayed a strong decrease in thickness, we set out to determine the presence of CUX1, a marker for the CP and future layer I-IV (Cubelos et al., 2008). CUX1 cells can be detected in the upper layers of the WT and *Tcf*4 heterozygous cortex, but is completely absent in the upper layers of the full *Tcf4* mutant cortex (**Figure 4G** CP). Interestingly, CUX1 is still present in the VZ and SVZ in all genotypes (**Figure 4G** VZ). Taken together, these data further substantiate that cortical differentiation and layer specification is affected upon loss of *Tcf4* and that this effect aggravates during development, leading to an absence of CUX1^+^ neurons in upper layers of the cortex in full *Tcf4* mutants.

### 3.5 Loss of *Tcf4* leads to defective development of the corpus callosum and hippocampus at E17.5

As shown above, deletion of *Tcf4* has a measurable effect on the neuronal differentiation within the developing cerebral cortex. To determine whether this effect is represented by axonal projections and development of cortex-derived structures, we performed GAP43 immunohistochemistry in combination with DAPI to visualize the axonal tracts in the cortex, hippocampus and the corpus callosum at E17.5 (Benowitz and Routtenberg, 1997; Grasselli and Strata, 2013; Strittmatter et al., 1995). GAP43 expression in the cortex can be divided at this stage in three compartments, an upper (CP), middle (layer VI), and lower (IZ) part, based on neuronal and axonal density. Quantification of the thickness of the compartments compared to the total thickness, shows a small but significant change in thickness of the middle compartment between the mutant- and both the WT (n=4; p<0.01, two-tailed) and heterozygous (n=4; p<0.01, two-tailed) animals of ~6% (**Figure 5A**).

As described above, we have detected a significant higher proportion of SATB2-CTIP2 double-positive neurons in full *Tcf4* mutants. Since SATB2-expressing cells are known to project through the corpus callosum (Alcamo et al., 2008; Leone et al., 2015; Srinivasan et al., 2012; Srivatsa et al., 2014), we set out to determine whether this potential mis-specification influences the development of the corpus callosum (**Figure 5B**). In both WT (**Figure 5B-1**) and heterozygous animals (**Figure 5B-2**) the corpus callosum seems unaffected (n=3), with the callosal bundle visualized through GAP43 IMHC and the callosal wedge (**Figure 5B** white arrowheads) by DAPI staining. Importantly, in the full *Tcf4* mutant (**Figure 5B-3**) the callosal bundle is completely absent (n=3; p<0.001, one-tailed), although the callosal wedge (**Figure 5B** white arrowheads) seems unaffected.

**Figure 5:**
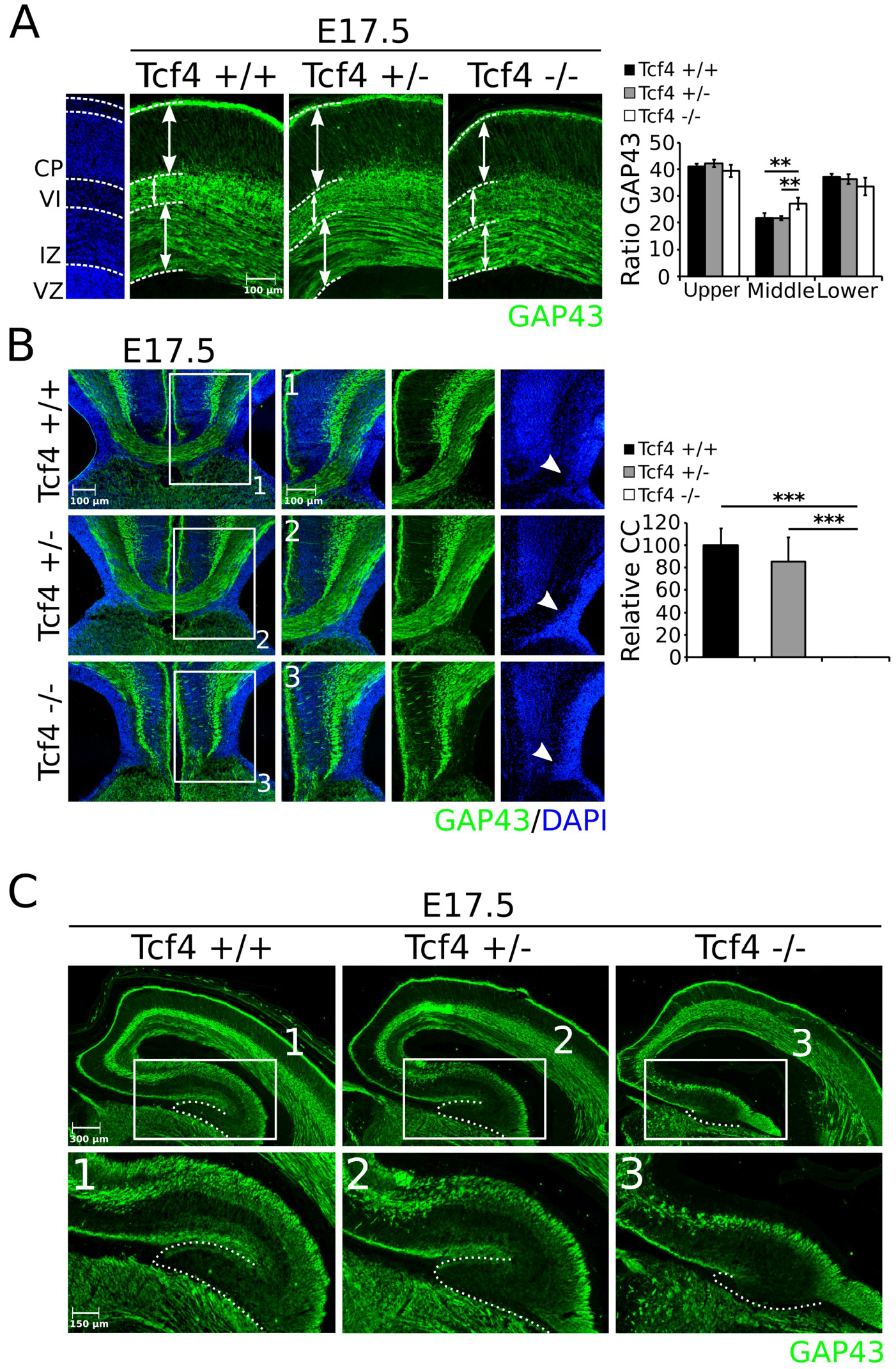
The *Tcf4* mutant shows an altered distribution of GAP43 in the cortex, a loss of the corpus callosum, and underdeveloped hippocampi at El7.5. (**A**) Distribution of GAP43 (green) is altered in the cortex of the *Tcf4* mutant at E17.5. Although the thickness of the upper (CP) and lower (IZ) compartments does not show a difference between WT (black bars), heterozygous (gray bars), and mutant (white bars) embryos, the thickness of the middle (VI) compartment is increased with ~5.4% in the mutant compared to WT (n=4; ** p<0.01, two-tailed) and heterozygous (n=4; ** p<0.01, two-tailed) embryos. Ratio to total thickness GAP43. (B) Development of the corpus callosum is strongly affected in the *Tcf4* mutant at E17.5. The callosal bundle, visualized by GAP43 (green), develops normally in the WT (**1**) and heterozygous (**2**) embryos, whereas it is lost in the mutant (**3**). The callosal wedge, visualized by DAPI (blue), appears to be normally present in WT, heterozygous, and mutant embryos (white arrowheads). Quantification of the callosal thickness shows a significant loss of ~100% between WT (black bar) and mutant (white bar) (n=4; *** p<0.001, one-tailed) and between heterozygous (gray bar) and mutant (n=4; *** p<0.001, one-tailed) embryos. WT was set at 100%. (**C**) The hippocampus is underdeveloped in the *Tcf4* mutant embryos at E17.5 compared to WT and heterozygous embryos. The underdevelopment of the hippocampus can mainly be detected in the dentate gyrus (**1-3**).

Examination of the hippocampus in the full *Tcf4* mutant showed an absence of the dentate gyrus (**Figure 5C-1** to **3** for magnifications), next to a clear incomplete development in heterozygous animals. Together, these data indicate that not only the development of the cerebral cortex is disrupted by the loss of *Tcf4*, also the development of cortex-related structures, like the corpus callosum and hippocampus, is affected at this stage of development.

### 3.6 Corpus callosum and hippocampal defects persists towards birth in *Tcf4* mutants

As shown above, the corpus callosum and hippocampus show an aberrant development upon loss of *Tcf4*. To determine whether this effect persists towards birth, we tracked the development of these structures by GAP43 IMHC at P0, similar as shown for E17.5 above (**Figure 6**). Whereas at E17.5 distribution of GAP43 only showed a significant change in the middle compartment, at P0 GAP43 distribution in the cerebral cortex displays mild but significant changes in all subsections analyzed (**Figure 6A**). The upper (CP) compartment of GAP43 is increased in both the heterozygous with ~3% (n=3 p<0.05, two-tailed) and mutant with ~9% (n=3 p<0.001, two-tailed) compared to WT. The difference of ~6% between the heterozygous and mutant animals is similarly significant (n=3 p<0.01, two-tailed). Interestingly, whereas at E17.5 the middle (V-VI) compartment was increased in thickness in the mutant, at P0 this compartment shows a decrease of ~6% between the WT and full mutants (n=3 p<0.05, two-tailed). In the lower (SP-IZ) compartment only a significant difference was detected between the WT and heterozygous animals of ~2.5% (n=3 p<0.05, two-tailed). The difference in GAP43 between E17.5 and P0 indicates that axonal tract formation in the cortical layers is affected even at late stages of development.

**Figure 6:**
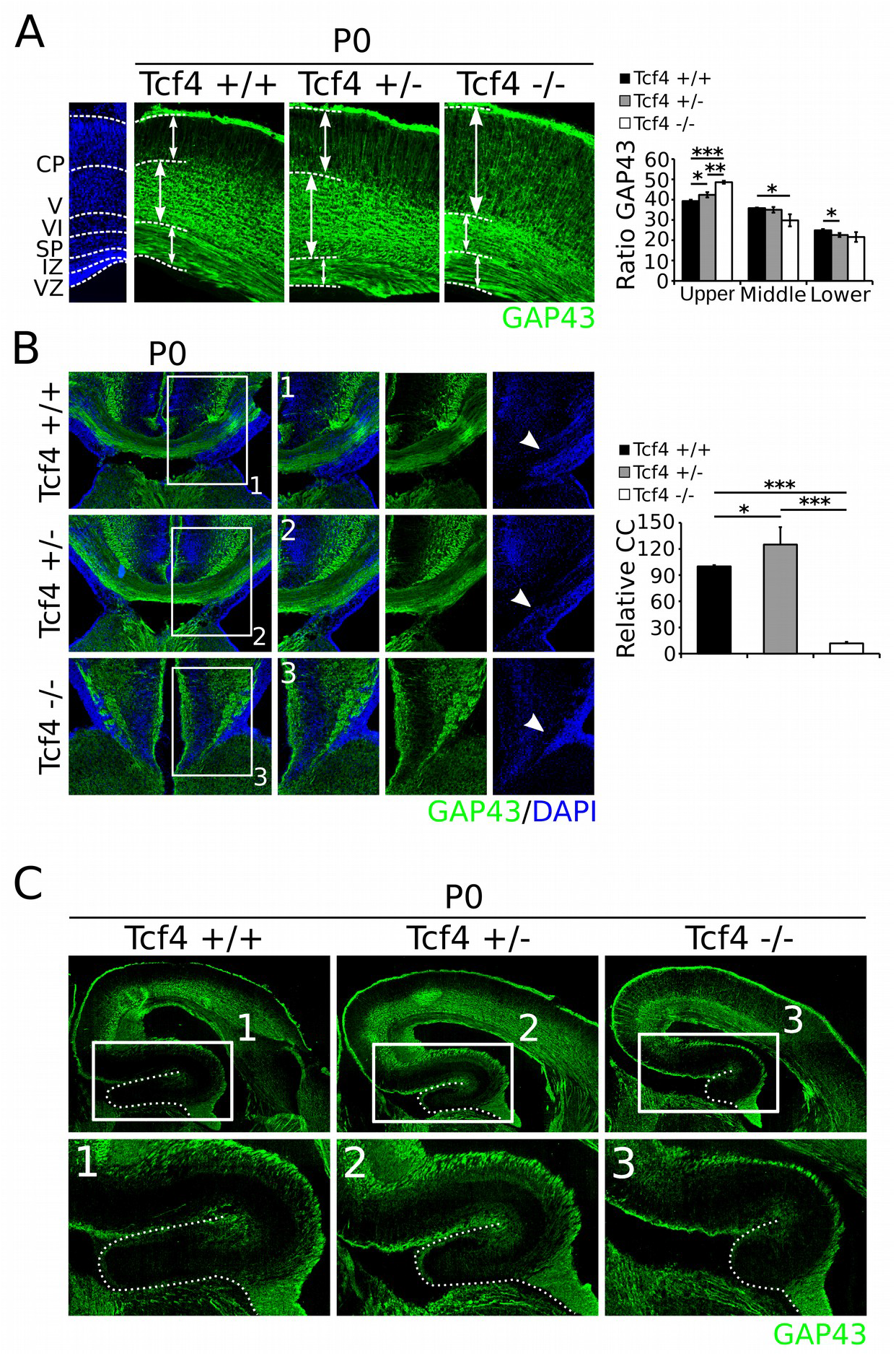
The *Tcf4* mutant shows an altered distribution of GAP43 in the cortex, a loss of the corpus callosum, and underdeveloped hippocampi at P0. (**A**) Distribution of GAP43 (green) in the cortex is altered in both heterozygous and mutant compared to WT animals at P0. Thickness of the upper (CP) compartment is increased with ~9% in the mutant compared to WT (n=3; *** p<0.001, two-tailed) and with ~6% compared to heterozygous animals (n=3; ** p<0.01, two-tailed). Heterozygous animals have an ~3% thicker upper compartment compared to WT animals (n=3; p<0.05, two-tailed). Thickness of the middle (V-VI) compartment is decreased with ~6% in the mutant animals compared to WT animals (n=3; p<0.05, two-tailed), whereas thickness of the lower (SP-IZ) compartment is not significantly altered between mutant and WT animals, but is decreased with ~3% in heterozygous compared to WT animals (n=3; * p<0.05, GAP43. (**B**)Ratio to total thickness GAP43. (**B**) Development of the corpus callosum is strongly affected in the *Tcf4* mutant at P0. The callosal bundle, visualized by GAP43 (green), develops normally in the WT (**1**), whereas it is increased in thickness in heterozygous (**2**) embryos, and is lost in the mutant (**3**). The callosal wedge, visualized by DAPI (blue), appears to be normally present in WT, heterozygous, and mutant embryos (white arrowheads). Quantification of the callosal thickness shows a significant loss of ~89% between WT (black bar) and mutant (white bar) (n=3; *** p<0.001, one-tailed) and ~114% between heterozygous (gray bar) and mutant (n=3; *** p<0.001, one-tailed) embryos. The corpus callosum shows an increase of ~25% in thickness in heterozygous animals compared to the WT (n=3; * p<0.05, one-tailed). WT was set at 100%. (**C**) The hippocampus is underdeveloped in the *Tcf4* mutant at P0 compared to WT and heterozygous animals. In the heterozygous animals the hippocampus appears to show a slight underdevelopment compared to the WT. The underdevelopment of the hippocampus can mainly be detected in the dentate gyrus (**1-3**).

The development of the corpus callosum is affected in full mutants (**Figure 6B-3**) compared to both WT (**Figure 6B-1**) (n=3 p<0.001, one-tailed) and heterozygous (**Figure 6B-2**) (n=3 p<0.001, one-tailed) animals at P0, confirming the initial defects observed at E17.5 including the apparent absence of a defective callosal wedge (**Figure 6B-1** to **3** white arrowheads). Confirming the data observed at E17.5, the development of the hippocampus, and specifically the dentate gyrus, is affected in full *Tcf4* mutants, although at this stage some initial cortical folding could be detected (**Figure 6C-1** to **3**). Taken together, the initial aberrations in cortical development, as identified at E17.5, persisted towards birth.

### 3.7 *Tcf4* acts as a transcriptional activator during cortical development and regulates genes involved in neuronal differentiation and maturation

In order to provide a better insight in the molecular mechanism of TCF4 action during cortical development we aimed to determine the early and therefore possible direct effects of *Tcf4* ablation through RNA-sequencing of E14.5 dissected cortices of WT and mutant embryos (n=3; 2 pooled embryos per biological replicate). These analyzes showed that *Tcf4* mainly acts as a transcriptional activator since the we observed 131 downregulated genes and 6 upregulated. Analysis of the top 25 upregulated (**Figure 7A**) and the top 5 downregulated genes (**Figure 7B**), based on fold-change of the transcript compared to WT samples indicates that *Tcf4* may regulate genes that are involved in the regulation of neuronal differentiation en neuronal migration (e.g. *NeuroD1, Mash1, Nos1*, and *Id2*) (Casarosa et al., 1999; Kim, 2013; Park et al., 2013; Zhang et al., 2015) and regulates its own expression (**Figure 7C**). Interestingly, these developmental processes that are typically affected in patients with ID. Confirming the data previously shown and described above, GO-term analysis (PANTHER Over-representation Test, **Figure 8A**) shows that *Tcf4* is involved in the regulation of (neuronal) differentiation, cell signaling, synaptic plasticity, and development of the telencephalon and hippocampus. Further analysis of the genes regulated by *Tcf4* shows that 26 of the 137 genes regulated by *Tcf4* (19%) have previously been shown to be mutated in cases of ID (**Figure 8B**) (Alders et al., 2014; Backx et al., 2010; Brockschmidt et al., 2007; Cocchella et al., 2010; Ehmke et al., 2017; Gerber et al., 2016; Gillentine et al., 2017; Guo et al., 2016; Labonne et al., 2016; Magoulas and El-Hattab, 2012; Merla et al., 2002; Metsu et al., 2014; Mikhail et al., 2011; Montesinos, 2014; Moore et al., 2016; Mulatinho et al., 2012; Myers et al., 2012; Nesbitt et al., 2015; Poot et al., 2010; Schoonjans et al., 2016; Schuurs-Hoeijmakers et al., 2013; Srivatsa et al., 2014; Tassano et al., 2015; Thevenon et al., 2014; Ţuţulan-Cunită et al., 2012). Characteristics of these cases range from moderate to severe ID, autistic phenotypes, brain malformations, speech impairments, and epilepsy. These traits can similarly be detected in patients with PTHS (Blake et al., 2010; Hasi et al., 2011; Peippo and Ignatius, 2012). Taken together, the transcriptome analyzes shows that *Tcf4* acts mainly as a transcriptional activator of the neurogenic profile and the genetic program clearly relates to neurodevelopmental disorders as PTHS and ID.

**Figure 7:**
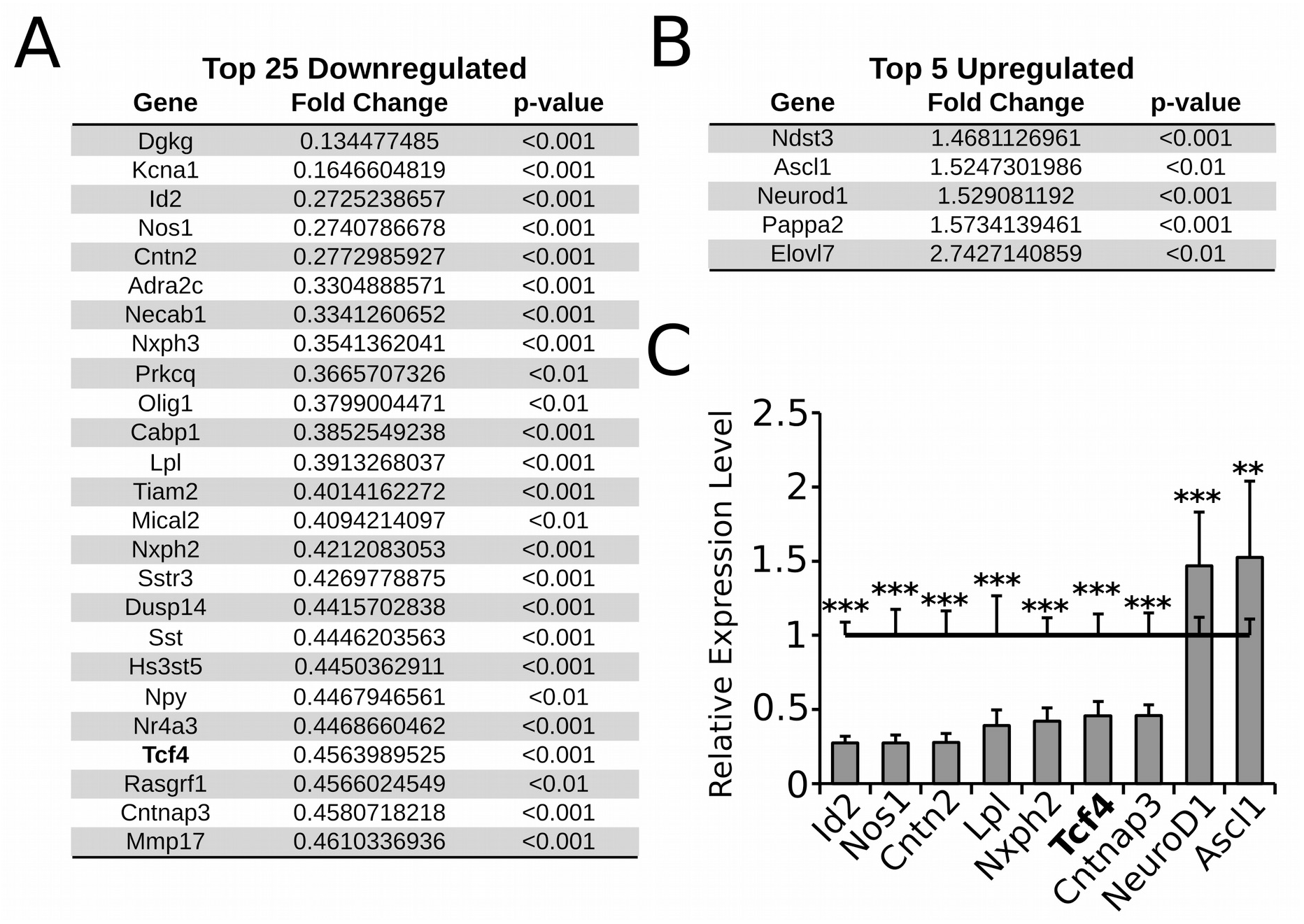
*Tcf4* top 25 up- and top 5 downregulated genes. (**A**) The top 25 downregulated genes in the *Tcf4* mutant based on significance and fold-change compared to WT samples. (**B**) The top 5 upregulated genes in the *Tcf4* mutant based on significance and fold-change compared to WT samples. (**C**) *Tcf4* is found to strongly regulate the expression of its own rest-transcript in the mutant. Many genes in the top-list are known disease-related genes, depicted in the graph.

**Figure 8:**
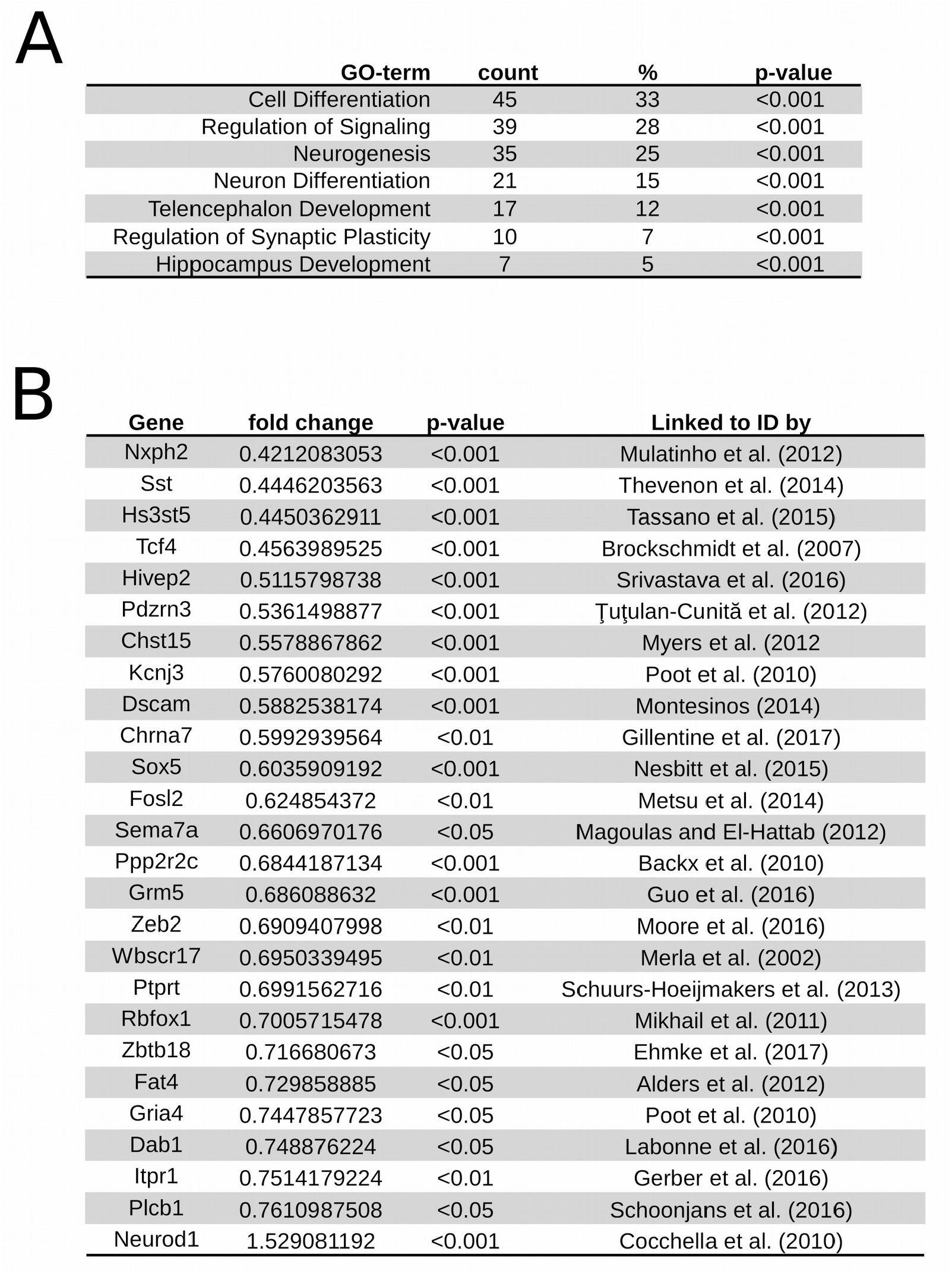
*Tcf4* is a transcriptional activator in E14.5 murine cortex and mainly regulates genes known to be mutated in cases of ID. (**A**) PANTHER Over-representation tests shows that *Tcf4* mainly regulates genes involved in neuronal differentiation, synapse regulation and cellular maturation at E14.5. Genes involved in the development of the telencephalon and hippocampus are similarly over-represented. (**B**) *Tcf4* regulates the expression of genes previously shown to be mutated in cases of ID. Table depicts gene name, fold change in the *Tcf4* E14.5 mutant cortex compared to the E14.5 WT cortex, p-value, and gene-specific references linking these genes to cases of ID.

## 4. Discussion

Here, we have shown that *Tcf4* is specifically expressed in different layers of the murine cortex during development and that it has a central role in cortical differentiation and maturation. Loss of *Tcf4* results in a disorganized cortex, in which the clear distinction between the different layers is merely affected, and postmitotic developing neurons are mis-specified. Furthermore, cortex-related structures, like the corpus callosum and hippocampus show clear developmental defects. Transcriptome analyzes through RNA-sequencing suggested that *Tcf4* acts mainly as a transcriptional activator involved in neuronal differentiation and maturation, and that *Tcf4* directly or indirectly regulates genes that are known to be mutated in cases of ID.

In humans, mutations within the bHLH domain in one copy of the *Tcf4* gene results in PTHS (Brockschmidt et al., 2007; Marangi et al., 2011; Marangi et al., 2012; Pontual et al., 2009; Sweatt, 2013), a rare mental disorder that is characterized by severe mental retardation, breathing abnormalities, and distinctive facial features (Blake et al., 2010; Hasi et al., 2011; Peippo and Ignatius, 2012). Brain-defects seen in PTHS patients include a smaller corpus callosum, bulging caudate nuclei, underdeveloped hippocampi, enlarged ventricles, and in some cases microcephaly (Blake et al., 2010; Ghosh et al., 2012; Hasi et al., 2011; Peippo and Ignatius, 2012). Many of these traits we were able to trace back in *Tcf4* mouse mutants, like the defects in the corpus callosum and hippocampi, initial underdevelopment of the cortex, and in some embryos we detected bulging caudate nuclei (**Supplemental Figure 2A**) or enlarged ventricles (**Supplemental Figure 2B**). In accordance to humans, loss of one copy of the *Tcf4* allele resulted in developmental defects, although we did detect a dose dependency, where the full *Tcf4* mutants displays the most consistent phenotypic characteristics as observed in human PTHS patients. The smaller hippocampi, initial underdevelopment of the cortex, and agenesis of the corpus callosum detected in this study match the data from Jung et al. (2018). However, when analyzing these mutants it is important to keep in mind that the study of Jung et al. focuses on the heterozygous mice of a different mouse model, which lacks exon 4 of the *Tcf4* gene, than used in this study, in which the bHLH domain of the *Tcf4* gene is replaced by a Neo-casette (Zhuang et al., 1996). As stated above patients with PTHS mainly show missense mutations or deletions in the basic region of the bHLH domain, resulting in a defective gene and possible protein in the human brain, indicating that the model used in this study could be used as a proper tool to mimic the effects of the main defects detected in PTHS patients.

Furthermore, we described that *Tcf4* is involved in the regulation of genes that are known to be mutated in cases of ID, together with traits like epilepsy, loss of the corpus callosum, and speech impairment also detected in patients with PTHS. For example patients with a mutation in *NeuroD1* show ID and speech impairments (Cocchella et al., 2010), whereas mutations in and *Zeb2*, related to the Mowat Wilson syndrome, is characterized by facial features, mental retardation, and an absent corpus callosum (Moore et al., 2016). Interestingly, although *Ascl1* is not directly related to ID, mutation of this gene is related to congenital central hypoventilation syndrome, a syndrome characterized by breathing abnormalities, which is also a known characteristic of PTHS patients (de Pontual et al., 2003). Besides its role in PTHS and other syndromes related to PTHS and ID, the *Tcf4* mouse mutant could be used in to better understand the onset and progression of autism and schizophrenia, both linked to *Tcf4* mutations (Brzózka and Rossner, 2013; Cousijn et al., 2014; de Munnik et al., 2014), with regard to its role in correct cortical development and synaptic plasticity. For example, in our RNA-sequencing data we detected a ~60% down-regulation of the *Lpl* gene, which is located on the 8p21.3 locus part of the 8p21 locus, a known schizophrenia susceptibility locus (Blouin et al., 1998; Fallin et al., 2011), and shows a ~42% and ~32% downregulation of *Kcnj3* and *Grm5* respectively, which are both associated to schizophrenia (Matosin et al., 2015; Yamada et al., 2012).

Taken together, our data show that *Tcf4* is of major importance in the murine brain for proper development of the cortex. Furthermore, loss of *Tcf4* leads to similar effects on the brain as observed in PTHS patients, making the *Tcf4* mutant mouse-line a good model to study molecular mechanisms of brain development and specifically in relation to PTHS patients. Our data provides a clear overview on the effects of *Tcf4* on brain development, next to previous roles described in development of the pons (Flora et al., 2007), and gives new insights in possible treatment of specific traits like PTHS and genes that could be involved in the development of PTHS-like syndromes and ID. Finally, our data set of genes regulated by *Tcf4* provide new leads for clinical genetics and or clinical diagnostics in, until now, genetically undefined neurodevelopmental disorders in humans.

## Authors’ Contributions

Conceived and designed the experiments: SM, MPS. Performed the experiments: SM, RB. Analyzed the data: SM, MPS. Contributed reagents/materials/analysis tools: SM, MPS. Wrote the paper: SM, MPS.

---

**Supplemental Figure 1:**
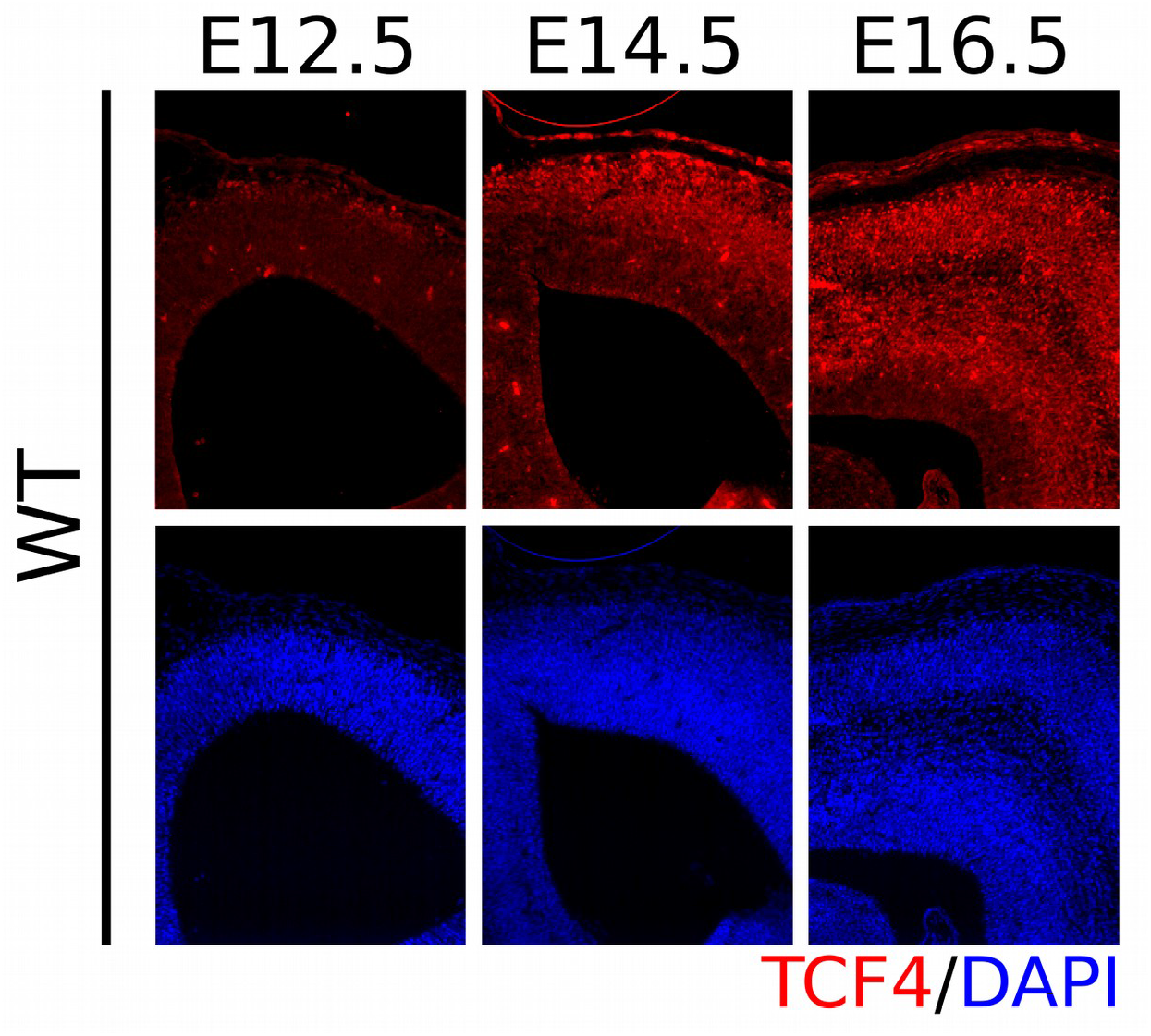
TCF4 is not clearly expressed in the murine cortex until E14.5. TCF4 expression is not detected in the murine cortex until E12.5 as shown by TCF4 immunohistochemistry (red) combined with DAPI (blue).

**Supplemental Figure 2:**
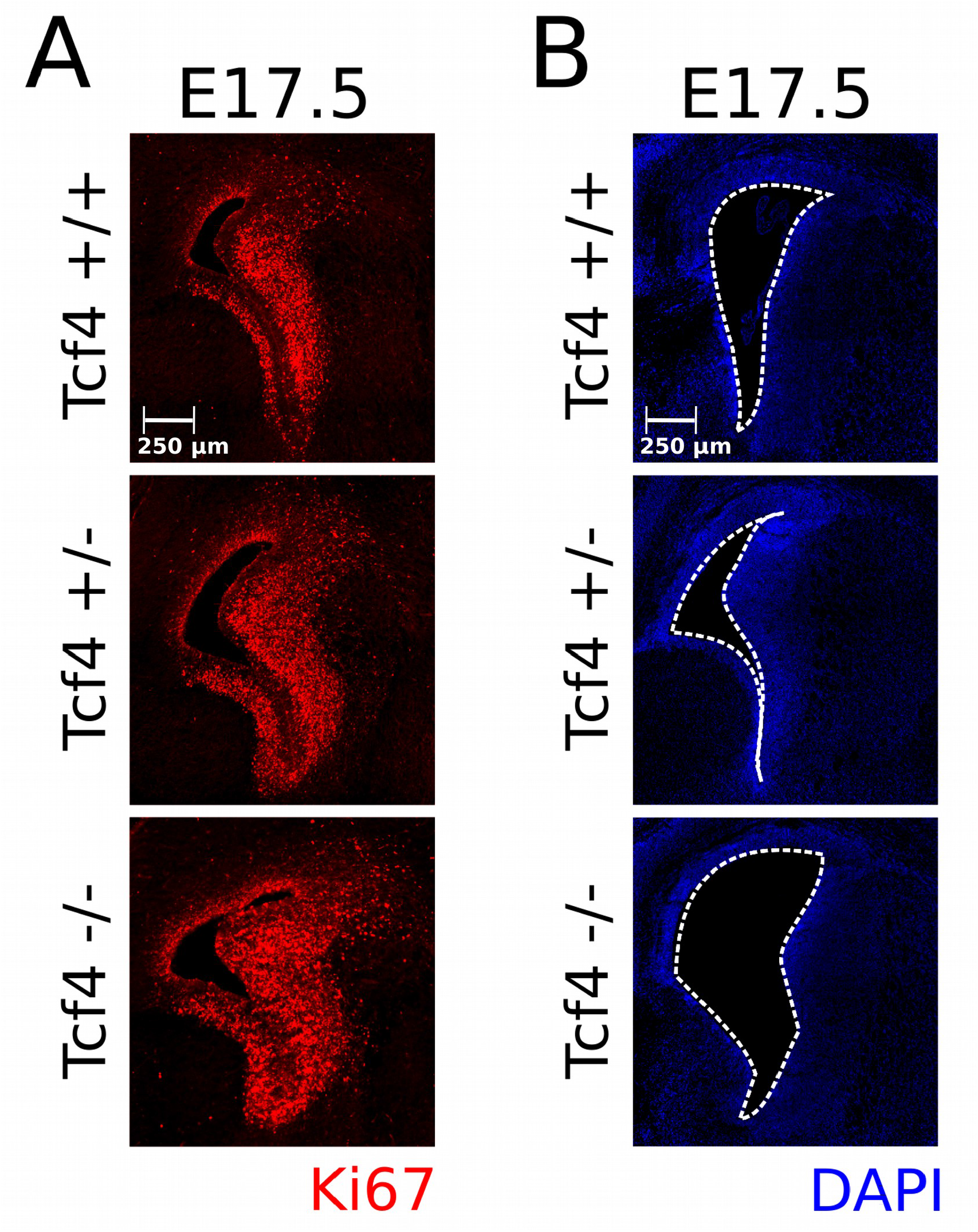
Bulging caudate nuclei and enlarged ventricles were detected in some of the *Tcf4* mutants at E17.5. (**A**) Bulging caudate nuclei were detected in some *Tcf4* mutant embryos at E17.5, which was accompanied by an increase in the presence of the proliferation marker Ki67 (red) staining. An intermediate phenotype was detected in some heterozygous littermates. (**B**) Similarly, enlarged ventricles were detected in some of the E17.5 *Tcf4* mutant embryos, visualized by DAPI (blue) staining.

